# Dissociable effects of surprising rewards on learning and memory

**DOI:** 10.1101/111070

**Authors:** N. Rouhani, K. A. Norman, Y. Niv

## Abstract

The extent to which rewards deviate from learned expectations is tracked by a signal known as a “reward prediction error”, but it is unclear how this signal interacts with episodic memory. Here, we investigated whether learning in a high-risk environment, with frequent large prediction errors, gives rise to higher fidelity memory traces than learning in a low-risk environment. In Experiment 1, we showed that higher magnitude prediction errors, positive or negative, improved recognition memory for trial-unique items. Participants also increased their learning rate after large prediction errors. In addition, there was an overall higher learning rate in the low-risk environment. Although unsigned prediction errors enhanced memory and increased learning rate, we did not find a relationship between learning rate and memory, suggesting that these two effects were due to separate underlying mechanisms. In Experiment 2, we replicated these results with a longer task that posed stronger memory demands and allowed for more learning. We also showed improved source and sequence memory for high-risk items. In Experiment 3, we controlled for the difficulty of learning in the two risk environments, again replicating the previous results. Moreover, equating the range of prediction errors in the two risk environments revealed that learning in a high-risk context enhanced episodic memory above and beyond the effect of prediction errors to individual items. In summary, our results across three studies showed that (absolute) prediction error magnitude boosted both episodic memory and incremental learning, but the two effects were not correlated, suggesting distinct underlying systems.

---

Reward prediction errors – phasic signals that track the difference between actual and expected outcomes – play a well-established role in updating stored information about the values of different choices in reinforcement learning. It is less clear, however, what role reward prediction errors play in the formation of episodic memories given that reinforcement learning and episodic memory have been traditionally studied separately.

Reward prediction errors are known to modulate dopamine release. Dopamine, in turn, modulates processing in the hippocampus, a key structure for episodic memory. This dopaminergic link therefore provides a potential mechanism for reward prediction errors to affect episodic memory. However, there are several ways by which reward prediction errors could potentially influence episodic memory. First, memory formation might be affected by the *signed* reward prediction error (i.e., expected minus actual reward); dopamine release tracks this signed prediction error, increasing when an experienced outcome is better than expected, and decreasing when the outcome is worse than expected (Schultz, Dayan, & Montague, 1997). If a signed prediction error signal influences episodic memory, we would expect a positive prediction error to improve memory whereas a negative prediction error would worsen it.

A second possibility is that the magnitude of the prediction error could influence episodic memory *regardless* of the sign of the error, enhancing memory for events that are either much better or much worse than expected. The effects of unsigned prediction errors are thought to be mediated by the locus-coeruleus-norepinephrine (LC-NE) system, which demonstrates a phasic response to unexpected changes in stimulus-reinforcement contingencies, regardless of sign, in both reward and fear learning (for a review, see Sara, 2009), and modulates increases in learning rate, i.e. the extent to which a learner updates their values, following large unsigned prediction errors (Behrens, Woolrich, Walton, & Rushworth, 2007; McGuire, Nassar, Gold, & Kable, 2014; Nassar et al., 2012). Importantly, recent evidence also indicates that the locus coeruleus co-releases dopamine with norepinephrine, giving rise to dopamine-dependent plasticity in the hippocampus (Kempadoo, Mosharov, Choi, Sulzer, & Kandel, 2016; Takeuchi et al., 2016). This latter pathway thereby provides a mechanism whereby unsigned prediction errors could affect episodic memory, by modulating hippocampal plasticity.

In our study, we set out to measure how both signed and unsigned reward prediction errors affect episodic memory formation. We also wanted to measure the effect of *risk context* (i.e., whether unsigned prediction errors were large or small, on average, in a particular environment) on episodic memory. Previous work on the effects of risk context show that dopamine signals scale to the reward variance of the learning environment (Tobler, Fiorillo, & Schultz, 2005), allowing for greater sensitivity to prediction errors in lower variance contexts. Moreover, behavioral learning rate and BOLD responses in the dopaminergic midbrain and striatum have been shown to reflect this adaptation, with higher learning rates and an increased striatal response to prediction errors when the reward variance is lower (Diederen, Spencer, Vestergaard, Fletcher, & Schultz, 2016). We therefore expected higher learning rates in a low-risk context, but it was unclear whether this effect would interact with episodic memory. If anything, we expected opposite effects, such that a high-risk context would induce better episodic memory, as increased arousal may lead to enhanced encoding of all items.

To investigate the effect of prediction errors and risk context on the structure of memory, we asked participants to learn by trial and error which of two types of images, indoor or outdoor scenes, led to higher rewards. Trial-unique indoor and outdoor images were presented in two different contexts or ‘rooms,’ with each room associated with a different degree of outcome variance (although indoor and outdoor images in the two rooms were matched in value, on average). At a later stage, memory for the scenes was assessed through recognition memory (‘item’ memory), identification of the room the item belonged to (‘source’ or context memory; Exp. 2-3), and the ordering of a pair of items (‘sequence’ memory). Choice trials further confirmed memory for the outcomes associated with the scenes.

## Experiment 1

In Experiment 1, we assessed whether reward prediction errors interact with episodic memory for rewarding events. Participants learned the values of two types of images (indoor or outdoor scenes) in two learning contexts (‘rooms’). The two learning contexts, while matched for mean value, had different degrees of reward variance (‘risk’) such that the rewards associated with images in the ‘high-risk room’ gave rise to higher absolute prediction errors than in the ‘low-risk room’.

### Method

#### Participants

Two hundred participants initiated an online task using Amazon Mechanical Turk (MTurk), and 174 completed the task. We obtained informed consent online, and participants had to correctly answer questions checking for their understanding of the instructions before proceeding; procedures were approved by Princeton University’s Institutional Review Board. Participants were excluded if they (1) had a memory score (*A’*: sensitivity index in signal detection; Pollack & Norman, 1964) of less than 0.5, or (2) missed more than three trials. These criteria led to the exclusion of ten participants, leading to a final sample of 164 participants.

#### Procedure

Participants learned by trial and error the value of two types of images (indoor or outdoor scenes) in two rooms defined by different background colors. In each room, one type of scene was worth 40¢ on average (low-value ‘scene’) and the other worth 60¢ (high-value ‘scene’). The average values of the scenes were matched across rooms, but the reward variance of the high-risk room was more than double that of the low-risk room (high-risk *σ* = 34.25, low-risk *σ* = 15.49). Participants were told that in each room one type of scene is worth more than the other (a ‘winning’ scene) and were asked to indicate the winner after viewing all images in a room. After the two learning blocks (one high-risk and one low-risk), participants completed a risk attitude questionnaire (DOSPERT, Weber, Blais, & Betz, 2002) that served to create a 5-10 minute delay between learning and memory tests. Participants then completed an item recognition task, after which they made choices between previously seen images.

##### Learning

On each trial, participants were shown a trial-unique image (either an indoor or outdoor scene) for 2 seconds. Participants were then given 5 seconds to estimate how much that type of scene is worth on average in that room (from 1 to 100 cents). The image was then presented again for 3 seconds along with its associated reward (see Figure 1A). Participants were told that their payment was not contingent on how accurate their guesses were, but instead was solely determined by the rewards they received (they were told that they would be paid a portion of the rewards they received, approximately ¼ of the actual outcomes). They were also instructed to pay attention to the images themselves as they would get a chance to choose between them later on in the experiment. There were 16 trials in each room (8 outdoor and 8 indoor). Rewards were 20¢, 40¢, 80¢, 100¢ (twice each) for the high-risk–high-value scene, 0¢, 20¢, 60¢, 80¢ for the high-risk–low-value scene, 45¢, 55¢, 65¢, 75¢ for the low-risk–high-value scene and 25¢, 35¢, 45¢, 55¢ for the low-risk–low-value scene. An identical sequence of rewards was applied across participants within rooms, with the order of the rooms (and risk levels) randomized.

**Figure 1.**
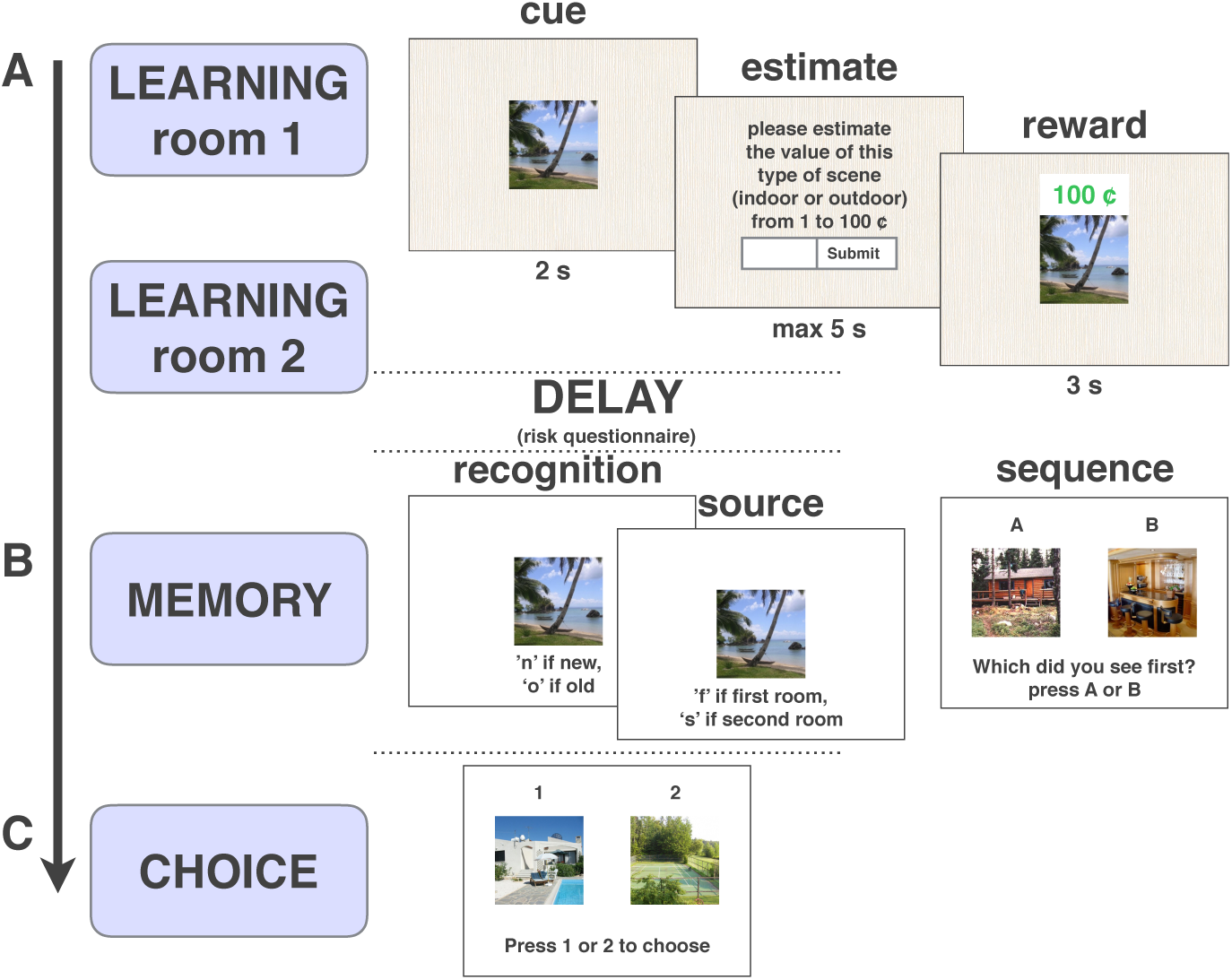
Task Design. A: Example learning trial. On each trial, participants were shown an image (“cue”), and were asked to estimate how much on average that type of scene (indoor or outdoor) was worth (“estimate”). They then saw the image again with a monetary outcome (“reward”). Each image appeared on one trial only. B: Memory tests. Participants completed item recognition, source (Exp. 2,3) and sequence memory tasks. C: Choice task. Participants chose between previously seen images that were matched for reward outcome, risk context, and/or scene value.

##### Memory

After completing the risk questionnaire, participants were presented with a surprise recognition memory test in which they were asked whether different scenes were old or new (see Figure 1B) as well as their confidence for that memory judgment (from 1 ‘guessing’ to 4 ‘completely certain’). There were 32 test trials, including 16 old images (8 from each room) and 16 foils. Participants were then asked to sequence 8 pairs of images (that were not included in the recognition memory test) by answering ‘which did you see first?’ (see Figure 1B) and by estimating how many trials apart the images had been from each other. Each pair belonged to either the low (4 pairs) or the high-risk room (4 pairs).

##### Choice

Finally, participants were asked to choose between pairs of previously seen images for another chance to receive their associated reward, thus assessing memory for their outcomes (see Figure 1C). The pairs varied in either belonging to the same room or different rooms and some were matched for reward and/or average scene value in order to determine which feature of reward learning gave rise to a choice preference. The choices were presented without feedback.

##### Statistical Analysis

Analyses were conducted using paired t-tests, repeated measures ANOVAs, and generalized linear mixed-effects models (R lme4 package, Bates et al., 2015). All results reported below (t-tests and ANOVAs) were confirmed using linear or generalized mixed-effects models treating subject as a random effect (for both the intercept and slope of the fixed effect in question).

### Results

#### Learning

Participants learned the average values of the high and low-value scenes better in the low-risk than in the high-risk room, as assessed by the deviation of their value estimates from the true averages of the scene values (*t*(163) = 14.52, p < 0.001; Figure 2A). Computing explicit prediction errors (i.e., the difference between the guess and the actual outcome) for the different rooms and scenes revealed that there were higher prediction errors in the high-risk room (*t*(163) = 36.77, p < 0.001; Figure 2B), as expected. Moreover, there was an interaction between risk and scene value such that participants overestimated the value of low-value scenes (resulting in negative prediction errors) and underestimated the value of high-value scenes (resulting in positive prediction errors) to a greater extent in the high-risk room than in the low-risk room, on average (*F*(1,163) = 141.2, p < 0.001; Figure 2C). This demonstrates more difficulty in separating the values of the scenes in the high-risk room, consistent with previous findings showing that when people estimate the means of two largely overlapping distributions, they tend to average across the two distributions, thereby grouping them into one category instead of separating them into two (Gershman & Niv, 2013).

**Figure 2.**
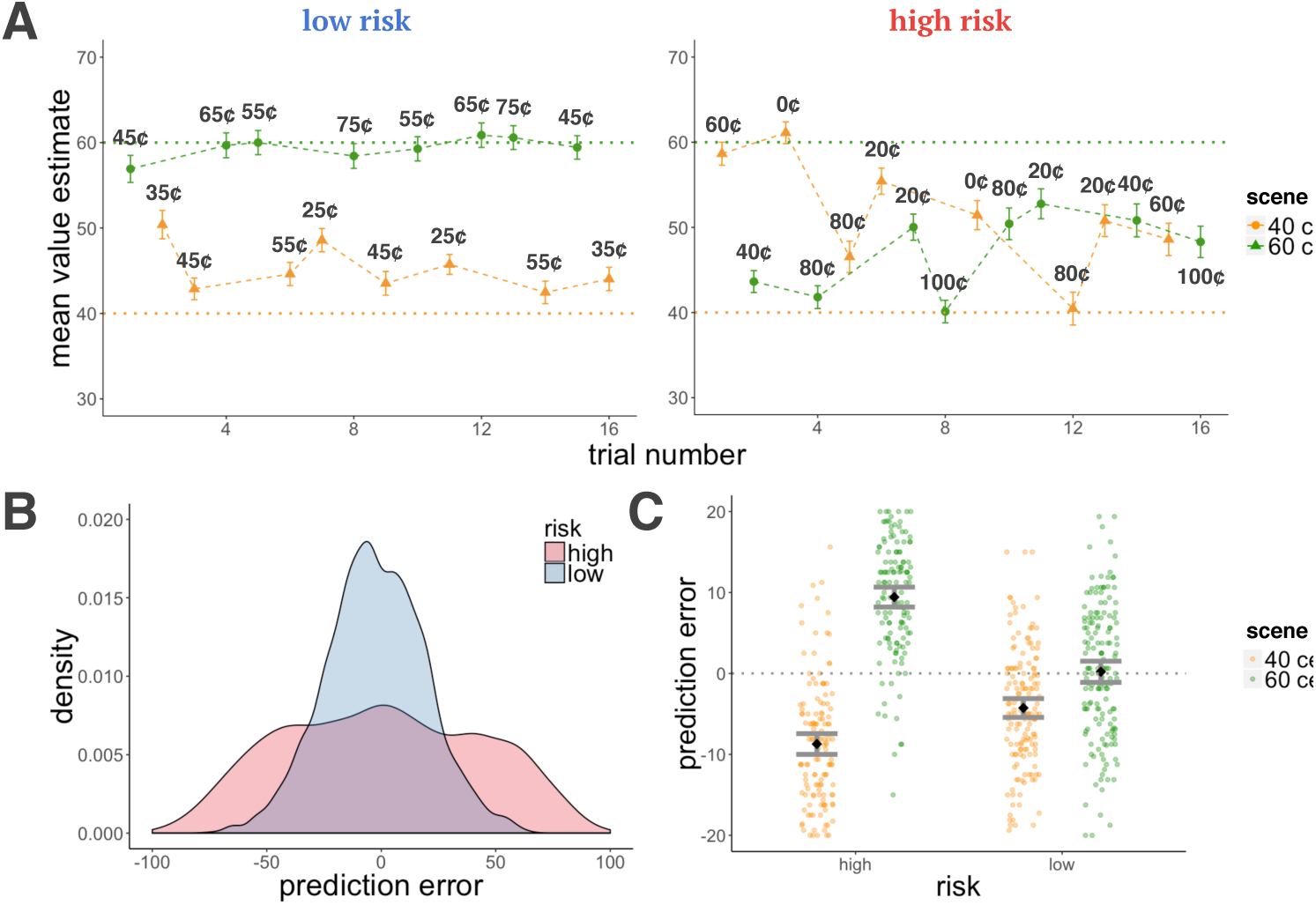
Experiment 1, learning results. A: Average estimates for the high and low-value scenes as a function of trial number for the high and low-risk rooms. Participants learned better in the low-risk room, indicated by the proximity of their guesses to the true values of the scenes (dashed horizontal lines). Cent values represent the outcome participants received on that trial (after entering their value estimate). B: Density plot of prediction errors in each room. There were more high-magnitude prediction errors in the high-risk in comparison to the low-risk room. C: There was an interaction for positive and negative prediction errors between risk context and scene value, such that participants overestimated the low-value scene and underestimated the high-value scene to a greater extent in the high-risk room. Error bars represent standard error of the mean.

#### Memory by Risk and Prediction Error

As predicted, high-risk items were remembered better than low-risk items (z = 2.37, p = 0.02, β = 0.31; Figure 3A). To test the effect of reward prediction errors on item memory, we first calculated trial-by-trial prediction errors by subtracting participants’ value estimates from the reward outcome they observed. We then used both signed and unsigned (absolute) prediction errors as regressors, together with a risk-level regressor in a mixed-effects logistic regression model of memory accuracy. We did not find signed prediction errors to influence memory (prediction error: z = 0.71, p = n.s., β = 0.04; risk: z = 2.29, p = 0.02, β = 0.30). Instead, we found that higher magnitude prediction errors enhanced memory regardless of the sign of the prediction error and explained the modulation of memory by risk (absolute prediction error: z = 3.36, p < 0.001, β = 0.23; risk: z = 0.9, p = n.s., β = 0.10; Figure 3B). This effect was significant also when controlling for reward outcome (absolute prediction error: z = 3.94, p < 0.001, β = 0.26; reward: z = 0.45, p = n.s., β = 0.02) and value estimates (absolute prediction error: z = 3.93, p < 0.001, β = 0.26; value: z = −0.09, p = n.s., β = −0.005), and when analyzing the high and low-risk items separately, (high: z = 1.90, p = 0.05, β = 0.18; low: z = 2.17, p = 0.03, β = 0.24). Reward prediction errors therefore affected episodic memory, such that larger deviations from one’s predictions, in any direction, enhanced encoding of items. Finally, there was no difference in sequence memory (the correct ordering of two images seen during learning) between pairs experienced in high and low-risk rooms (z = 0.11, p = n.s., β = 0.02).

**Figure 3.**
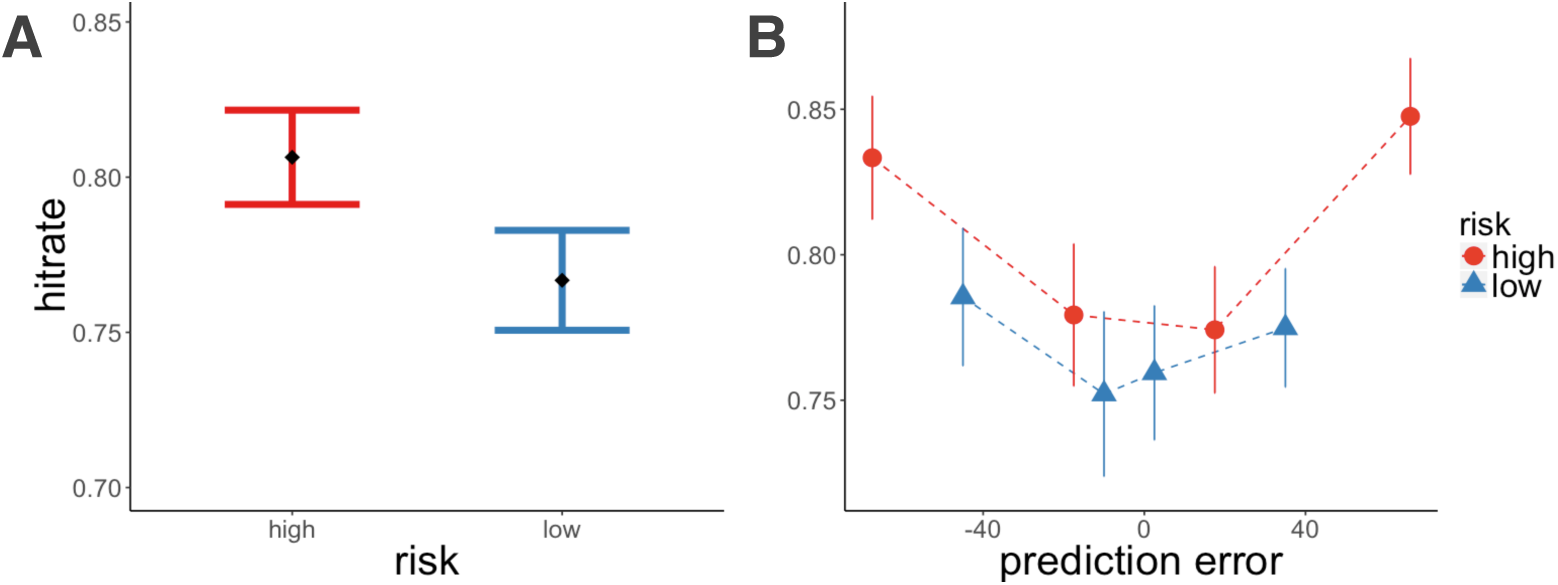
Experiment 1, recognition memory results. A: Recognition memory was better for high-risk items. B: There was better memory for high absolute prediction error items (controlling for risk context). Item memory was binned by the quartile values of prediction errors within each risk room. Each dot represents the average value within that quartile. Error bars represent standard error of the mean.

#### Learning Rate by Risk and Prediction Error

We also examined the effects of risk and prediction errors on the reward learning process itself. For this we calculated a trial-by-trial learning rate *α_t_* as the proportion of the current prediction error *R_t_ − V_t_* that was applied to update the value for the next encounter of the same type of scene, *V_t_*_+ 1_. Note that this is derived directly from the standard reinforcement learning update equation: *V_t_*_+1_ = *V_t_* + *α* _*t*_(*R_t_* – *V_t_*):

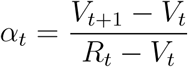

In agreement with recent findings (e.g.Diederen et al., 2016), we found that average learning rate was higher in the low-risk than in the high-risk room (*t*(163) = 3.37, p < 0.001; Figure 4A). Moreover, higher absolute prediction errors increased learning rates above and beyond the effect of risk in a mixed-effects linear model of learning rate (absolute prediction error: t = 3.30, p = 0.001, β = 0.07; risk: t = 4.67, p < 0.001, β = 0.16; Figure 4B). These results show that greater absolute prediction errors enhance value updating, and further that learning rates adapt to the reward variance of the context suggesting greater sensitivity to prediction errors in a lower risk environment. Controlling for absolute prediction error, we did not however find learning rate to predict memory on the current trial (learning rate: z = 0.85, p = n.s., β = 0.08; absolute prediction error: z = 3.42, p < 0.001, β = 0.20), nor the subsequent trial (learning rate: z = 0.56, p = n.s., β = 0.05; absolute prediction error: z = 3.06, p = 0.002, β = 0.19), demonstrating that learning rate increases were not correlated with memory even though both were enhanced by higher magnitude prediction errors.

**Figure 4.**
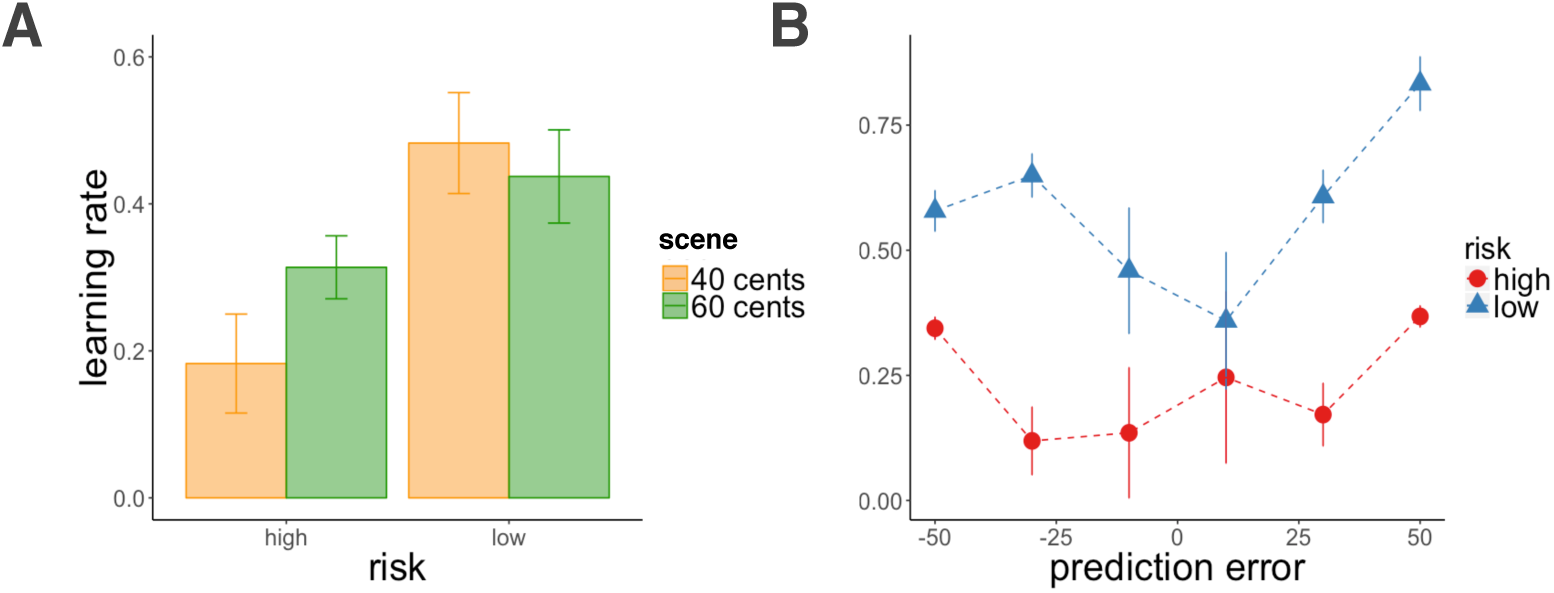
Experiment 1, learning rate results. A: Learning rate was higher in the low-risk context. Average learning rate plotted by risk context and scene value. B: Both absolute prediction errors and a low-risk context increased learning rate. Learning rates were binned by prediction errors (each dot represents the average prediction error within the binned range). Error bars represent standard error of the mean.

#### Choice by Reward and Value Difference

In the last test, participants were asked to make choices between pairs of previously-seen images that were matched for risk context, scene value, and/or reward outcome. We separately analyzed pairs that had matched reward outcomes and pairs that had unmatched outcomes. As the difference between the rewards previously associated with the two images increased, subjects chose the more rewarding image more often in a mixed-effects logistic regression model of choice (z = 6.40, p < 0.001, β = 0.54; Figure 5A). For choices with matched reward outcomes, in contrast, we could expect subjects to be indifferent between the two images. We instead found that subjects relied on their original value estimates for the two options in order to make their choices, such that the more they had valued one scene relative to the other, the more likely they were to choose it (z = 3.74, p < 0.001, β = 0.01; see Figure 5B). Importantly, the modulator of choice here was the initial guess of the value of the scene, rather than the outcome actually associated with the image (which was identical for the two images in question). We additionally found that, even when the two options had led to different rewards, the difference in initial value estimates for the scene was a significant predictor of choice, when controlling for the difference in actual reward outcome (value estimate difference: z = 2.27, p = 0.02, β = 0.16; reward difference: z = 7.25, p < 0.001, β = 0.52). We did not find risk level, the true average scene value, nor the difference in absolute prediction error between the images to additionally influence choice preference.

**Figure 5.**
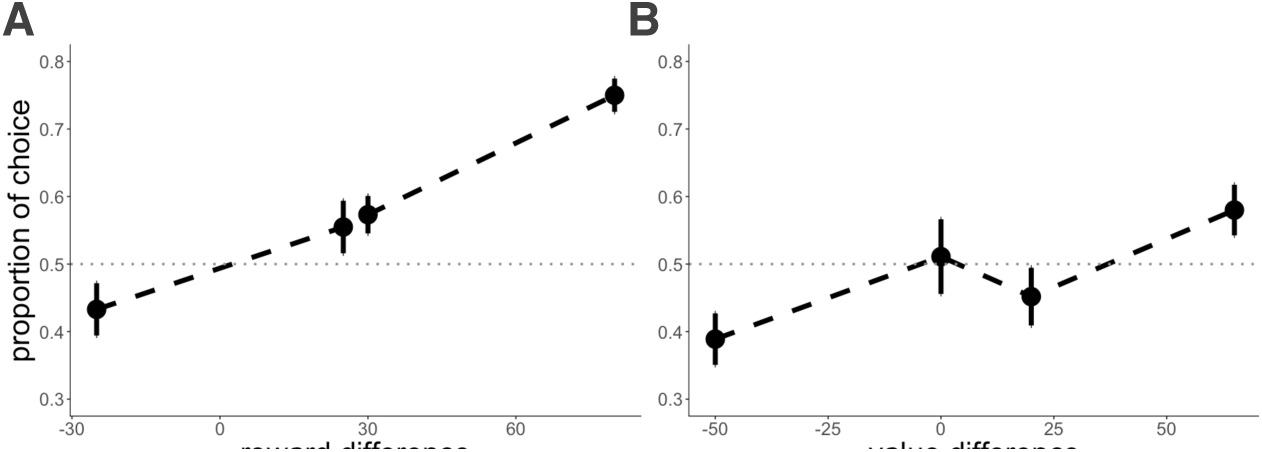
Experiment 1, choice results. A: For choices between options with different reward outcomes (each dot represents the actual reward difference) participants chose the image representing a larger reward. B: For choices between options with matched reward outcomes (each dot represents the average of the value differences binned by quartile), participants chose the image that they had valued more. Error bars represent the standard error of the mean.

## Experiment 2

Experiment 1 showed that deviations from predictions enhance episodic memory for the items that were associated with those predictions. We also demonstrated that both risk context and absolute prediction errors modulate learning rate. In particular, learning rate was higher in a low-risk environment, suggesting greater sensitivity to prediction errors during learning in this context, and further, in both contexts, high absolute prediction errors increased learning rate; however, we did not find learning rate to predict memory. Lastly, although we expected participants to select the more rewarding option when given a choice, we found that participants’ choices were also influenced by their own predictions of the value of that scene.

Notably, in contrast to standard reinforcement learning settings, our experiment involved only 16 trials of learning in each context, 8 for each ‘scene’. This initial learning phase is characterized by increased prediction errors and uncertainty relative to later learning, which might affect the relationship between prediction errors and episodic memory. Additionally, participants in Experiment 1 all experienced the same reward sequence, which inadvertently introduced regularities in the learning curves that could have also influenced initial learning and memory results. Finally, in this relatively short experiment, average memory performance was near ceiling (*A’* = 0.90). In Experiment 2, we therefore sought to replicate the results of Experiment 1 while increasing the number of learning and memory trials and randomizing reward sequence. With more trials, we were also able to test for sequence memory for items that were presented further apart in time, and we included a measure of source memory, a marker of episodic memory for the context of the probed item (i.e., which room the item belonged to).

### Method

#### Participants

Two hundred participants initiated an online task run on Amazon Mechanical Turk, and 148 completed the task. Following the same protocol as in Experiment 1, twelve subjects were excluded from the analysis leading to a final sample of 136 participants.

#### Procedure

The procedure was the same as in Experiment 1 but with some changes to learning, memory and choice. As in Experiment 1, rewards had a mean of 60¢ for the high-value scene and 40¢ for the low-value scene (high-risk–high-value scene: 20¢, 40¢, 60¢, 80¢, 100¢; high-risk–low-value scene: 0¢, 20¢, 40¢, 60¢, 80¢; low-risk-high-value scene: 40¢, 50¢, 60¢, 70¢, 80¢; low-risk–low-value scene: 20¢, 30¢, 40¢, 50¢, 60¢). We increased the number of learning trials from 16 to 30 trials per room, and we pseudo-randomized the reward sequence such that the rewards were drawn at random and were sampled three times without replacement.

During the item memory test, we asked participants to indicate whether items identified as ‘old’ belonged to the first or second room (see Figure 1B), as a measure of source memory. Additionally, given that sequence memory improves with greater distance between events (DuBrow & Davachi, 2013), we asked participants to order items that were further apart than in Experiment 1(13-14 trials apart). Finally, in the choice task, participants chose only between pairs matched for reward outcome.

### Results

#### Learning

As in Experiment 1, participants learned better in the low-risk than in the high-risk room (assessed by the average deviation of subject’s value estimates from the true means of the scene values; *t*(135) = 13.11, p < 0.001; Figure 6A). They experienced larger absolute prediction errors in the high-risk room (*t*(135) = 39.65, p < 0.001; Figure 6B), and there was again an interaction between risk and scene value such that subjects overestimated the value of the low-value scene, and underestimated the high-value scene to a greater extent in the high-risk room throughout learning (*F*(1,135) = 77.5, p < 0.001; Figure 6C).

**Figure 6.**
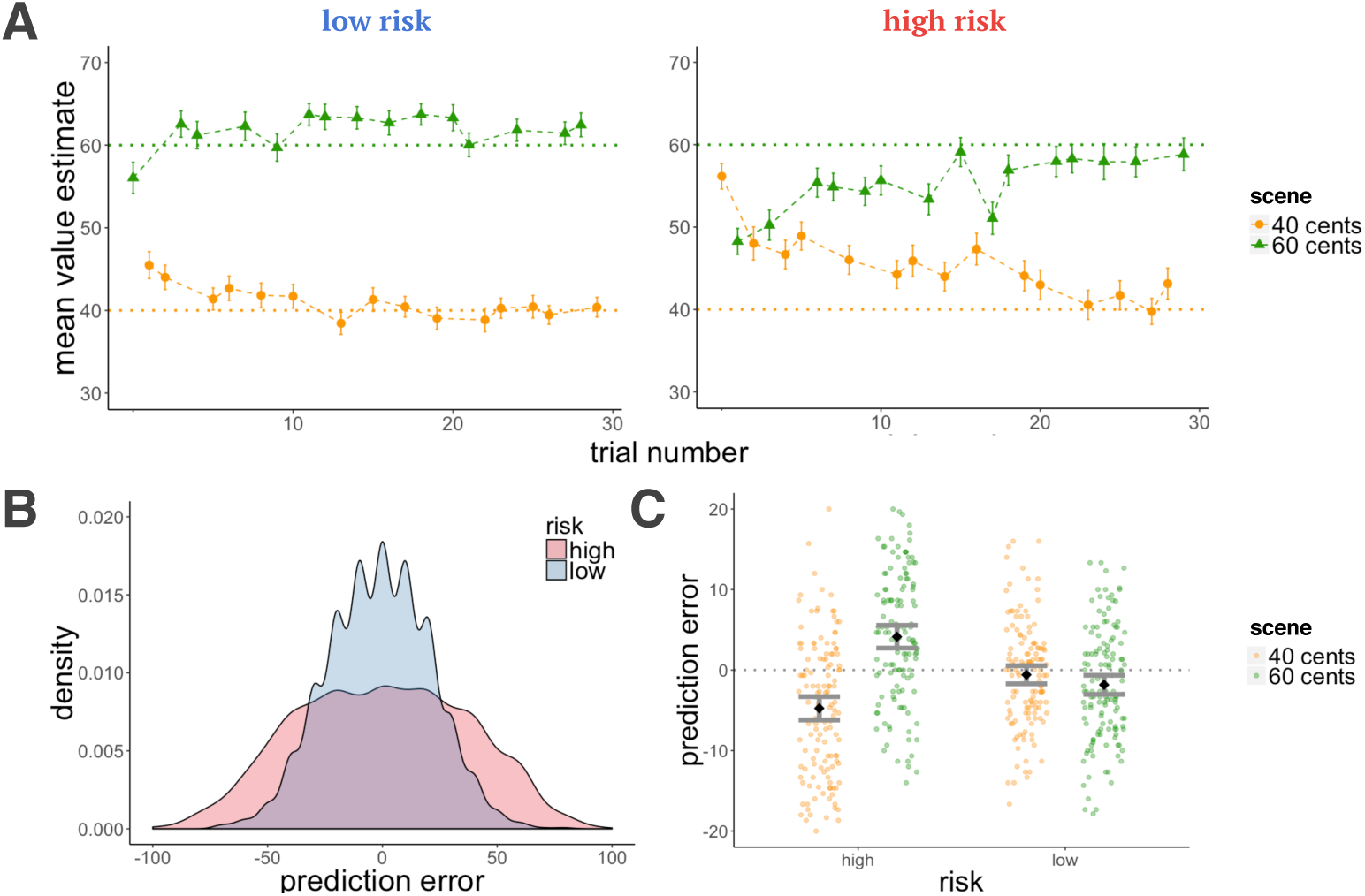
Experiment 2, learning results. A: Average estimates for the high and low-value scenes as a function of trial number for the high and low-risk rooms. Participants learned better in the low-risk room, indicated by the proximity of their guesses to the true values of the scenes (dashed horizontal lines). B: Density plot of prediction errors in each room. There were more high-magnitude prediction errors in the high-risk in comparison to the low-risk room. C: There was an interaction for positive and negative prediction errors between risk context and scene value, such that participants overestimated the low-value scene and underestimated the high-value scene to a greater extent in the high-risk room. Error bars represent standard error of the mean.

#### Memory by Risk and Prediction Error

By increasing the number of learning and memory trials, we significantly reduced average memory performance from Experiment 1 (*A’* = 0.86, *t*(275.23) = 3.04, p = 0.003). We nevertheless replicated the main results of Experiment 1: high-risk items were better remembered than low-risk items (z = 2.51, p = 0.01, β = 0.19; Figure 7A), and higher absolute prediction errors enhanced memory for those events (absolute prediction errors: z = 3.44, p < 0.001, β = 0.16; risk: z = 1.76, p = 0.08, β = 0.14, Figure 7B). Like in Experiment 1, this effect was significant when controlling for reward outcome (absolute prediction error: z = 4.14, p < 0.001, β = 0.18; reward: z = −1.71, p = n.s., β = −0.06) as well as for value estimates (absolute prediction error: z = 4.15, p < 0.001, β = 0.19; value: z = −1.16, p = n.s., β = −0.04).

**Figure 7.**
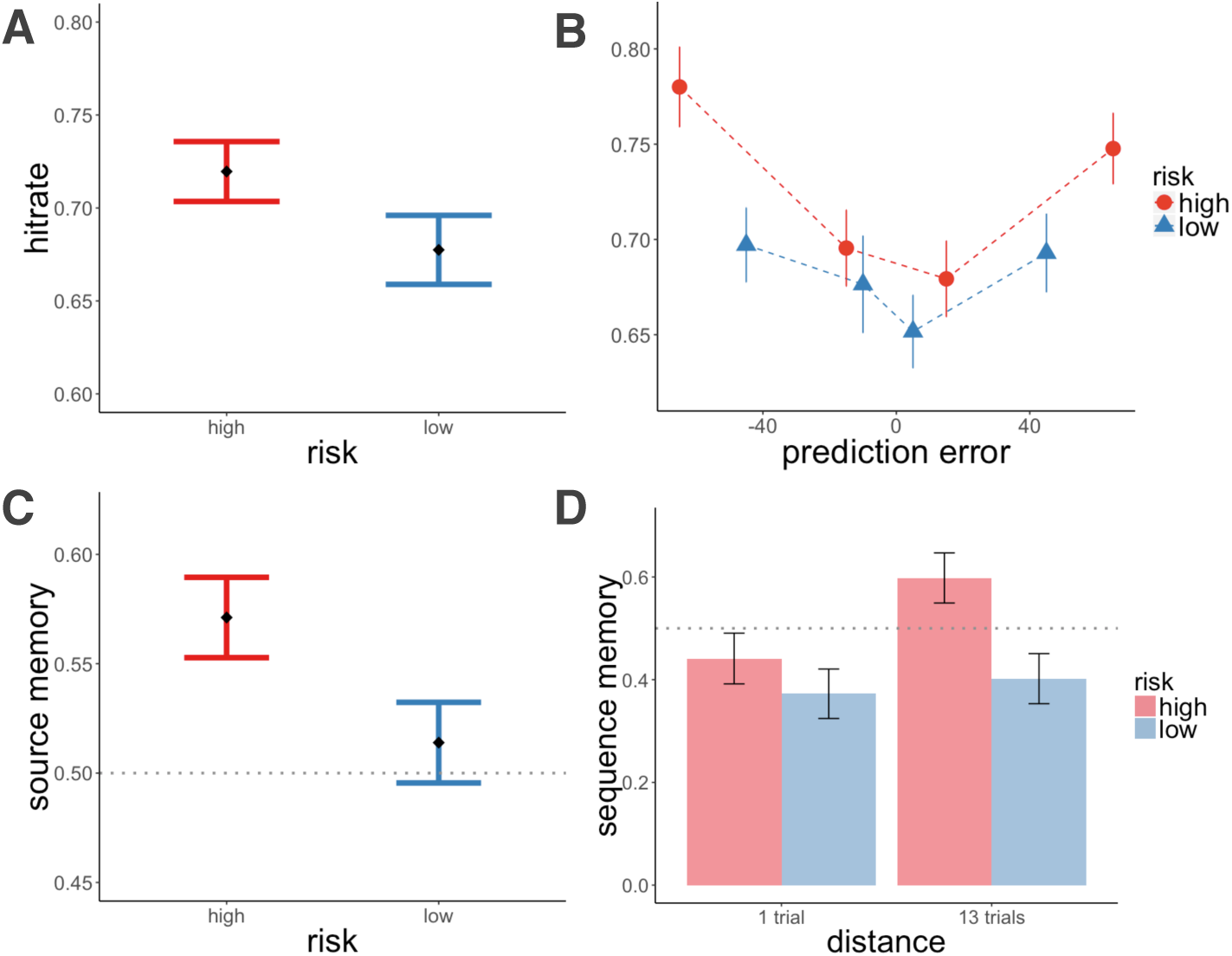
Experiment 2, memory results. A: Recognition memory was better for high-risk items. B: Absolute prediction errors enhanced item memory while controlling for risk context. Item memory was binned by the quartile values of prediction errors within each risk room, each dot represents the average value within that quartile. C: For correctly remembered items, source memory was better for high-risk items. D: A high-risk context and distance between items (number of trials between pairs) increased sequence memory. Error bars represent the standard error of the mean.

In addition, for the items correctly identified as old, we found better source memory for high-risk items (z = 2.05, p = 0.04, β = 0.25, Figure 7C). This effect was not modulated by absolute prediction error. Rather, it was a context effect: the source of a remembered image was better remembered if that item belonged to a high-risk context (absolute prediction errors: z = −0.60, p = n.s., β = −0.03; risk: z = 2.17, p = 0.03, β = 0.27). To verify that participants were not simply attributing ‘remembered’ items to the high-risk context, we looked at the proportion of high-risk source judgments for recognition hits and false alarms separately. We did not find a greater proportion of high-risk source judgments for false alarms, indicating that participants were not biased to report that ‘remembered’ items belonged to a high risk context (for high-risk hits: mean = 0.57, standard error = 0.02; for false alarms: mean = 0.49, standard error = 0.04; chance response is at 0.50). In this experiment, participants also exhibited better sequence memory for pairs from the high-risk context (z = 2.70, p = 0.007, β = 0.56, Figure 7D). Although we did not see this effect in Experiment 1, in Experiment 2 we had pairs that were separated by more trials (distant items were 13 and 14 trials apart). Indeed, greater distance predicted better sequence memory, controlling for risk (distance: z = 1.92, p = 0.05, β = 0.39; risk: z = 2.70, p = 0.006, β = 0.56). We therefore replicated our original results and further showed that other forms of episodic memory, source and sequence memory, were also enhanced in a high-risk context.

#### Learning Rate by Risk and Prediction Error

We replicated the results of Experiment 1 with respect to learning rates as well: participants had higher learning rates for the low-risk relative to the high-risk room, and higher absolute prediction errors additionally increased learning rates (absolute prediction errors: t = 5.12, p < 0.001, β = 0.09; risk: t = 7.01, p < 0.001, β = 0.18; Figure 8A-B). Controlling for absolute prediction error, we again did not find learning rate to predict memory on the current trial (learning rate: z = −0.29, p = n.s., β = −0.01; absolute prediction error: z = 4.44, p < 0.001, β = 0.20), nor the subsequent trial (learning rate: z = 0.68, p = n.s., β = 0.03; absolute prediction error: z = 3.53, p < 0.001, β = 0.17).

**Figure 8.**
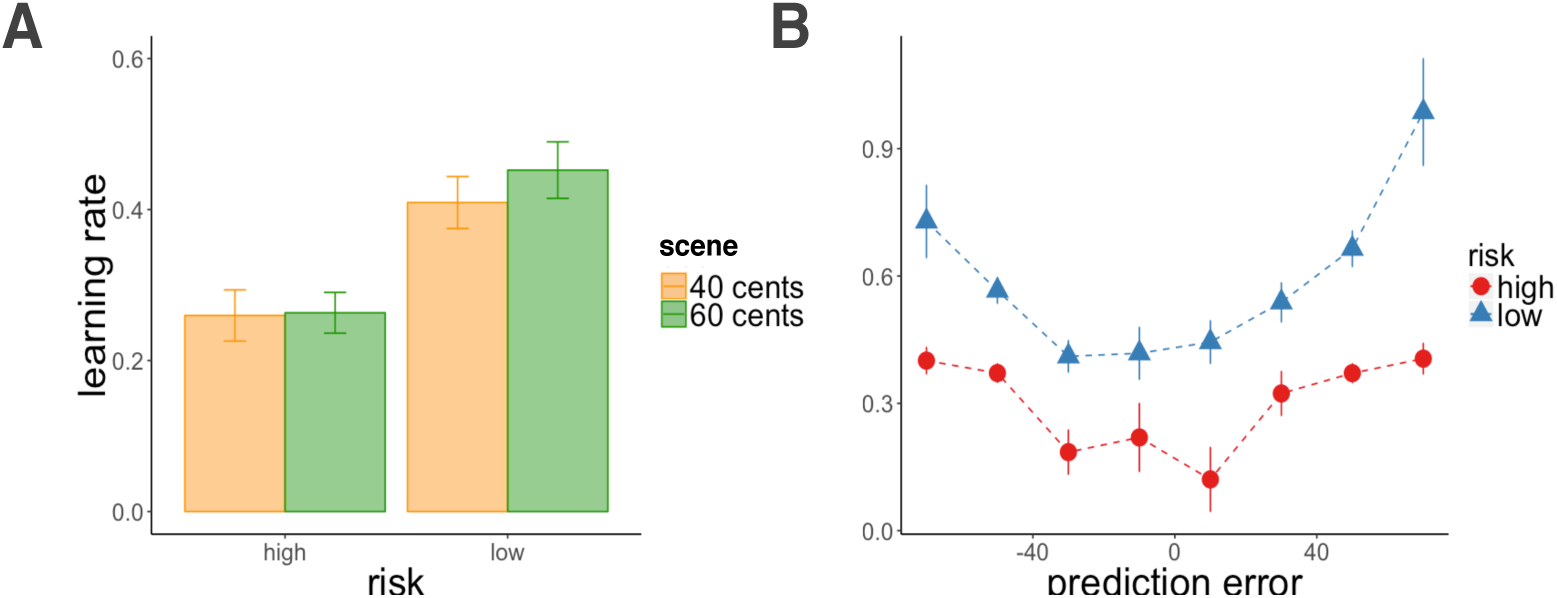
Experiment 2, learning rate results. A: Learning rate was higher in the low-risk context. Average learning rate plotted by risk context and scene value. B: Both absolute prediction errors and a low-risk context increased learning rate. Learning rates were binned by prediction errors (each dot represents the average prediction error within the binned range). Error bars represent standard error of the mean.

#### Choice by Value Difference

In this experiment, all choices were between images with matched reward outcomes. We replicated the results of Experiment 1 such that choice was predicted by the difference in subjects’ initial value estimates for the scenes (z = 2.78, p = 0.005, β = 0.18, Figure 9). In particular, even in this better-powered test (12 choice trials as compared to 4 choice trials with matched outcomes in Experiment 1), we did not see evidence for preference of images from one risk context versus the other (z = −1.56, p = n.s., β = −0.08).

**Figure 9.**
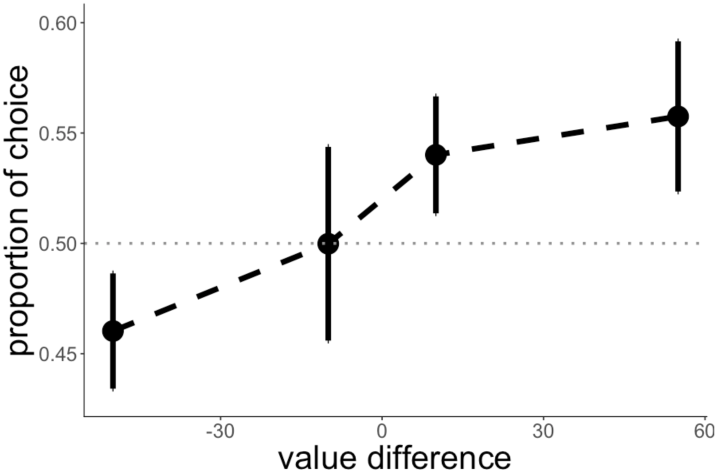
Experiment 2, choice results. For choices between options with matched reward outcomes, participants chose the scene that they had valued more. Each dot represents the average of the value differences binned by quartile. Error bars represent the standard error of the mean.

## Experiment 3

In Experiment 2, we doubled the number of training trials and replicated the results of Experiment 1 while extending the memory enhancement effects of high-risk learning to source and sequence memory. Nevertheless, a possible confound of the effects of risk on memory and learning in both experiments is that there was higher overlap between the outcomes for the indoor and outdoor scenes in the high-risk context as compared to the low-risk context. The distributions shared values from 20¢ to 80¢ (Exp. 1 & 2) in the high-risk room, but only 45¢ to 55¢ (Exp. 1) and 40¢ to 60¢ (Exp. 2) in the low-risk room. This greater overlap in the high-risk context could have made learning more difficult in comparison to the low-risk room, and therefore influenced the effects of absolute prediction error on subsequent memory. To test for this possibility, in Experiment 3 we made the learning conditions in the two rooms more similar by eliminating any overlap between the scene values.

### Method

#### Participants

We conducted a simulation-based power analysis on the effect of absolute prediction errors on item memory to achieve 80% power. This revealed that we would have sufficient power to replicate the results of Experiments 1 and 2 with as few as 55 participants. As a result, we had 100 participants initiate the study, of which 86 completed the task. Three participants were excluded based on our exclusion criteria (see Experiment 1) leaving a sample of 83 participants.

#### Procedure

We followed the same procedure as Experiment 2 but changed the rewards such that they had a mean of 80¢ for the high-value scene and 20¢ for the low-value scene, and there was no overlap between the rewards from the two scenes (high-risk–high-value scene: 60¢, 70¢, 80¢, 90¢, 100¢; high-risk–low-value scene: 0¢, 10¢, 20¢, 30¢, 40¢; low-risk-high-value scene: 70¢, 75¢, 80¢, 85¢, 90¢; low-risk–low-value scene: 10¢, 15¢, 20¢, 25¢, 30¢).

### Results

#### Learning

As in Experiment 1 and 2, participants learned better in the low-risk than in the high-risk room (*t*(82) = 6.28, p < 0.001; Figure 10A). However, learning in the two rooms was more similar here than in Experiment 2, as assessed by comparing subject learning differences between high and low risk contexts for Experiment 2 and 3 (*t*(148.98) = 1.84, p = 0.03; subject learning for each room was calculated by averaging trial-by-trial deviations of value estimates from the true means of the scene values). In contrast to the first two experiments, the range of prediction errors in the two rooms was also more similar in comparison to Experiment 1 and 2 (Figure 10B). As in previous experiments, there was an interaction between risk and scene value such that subjects overestimated the low-value scene and underestimated the high-value scene more in the high-risk than in the low-risk room, on average (*F*(1,82) = 23.02, p < 0.001; Figure 10C).

**Figure 10.**
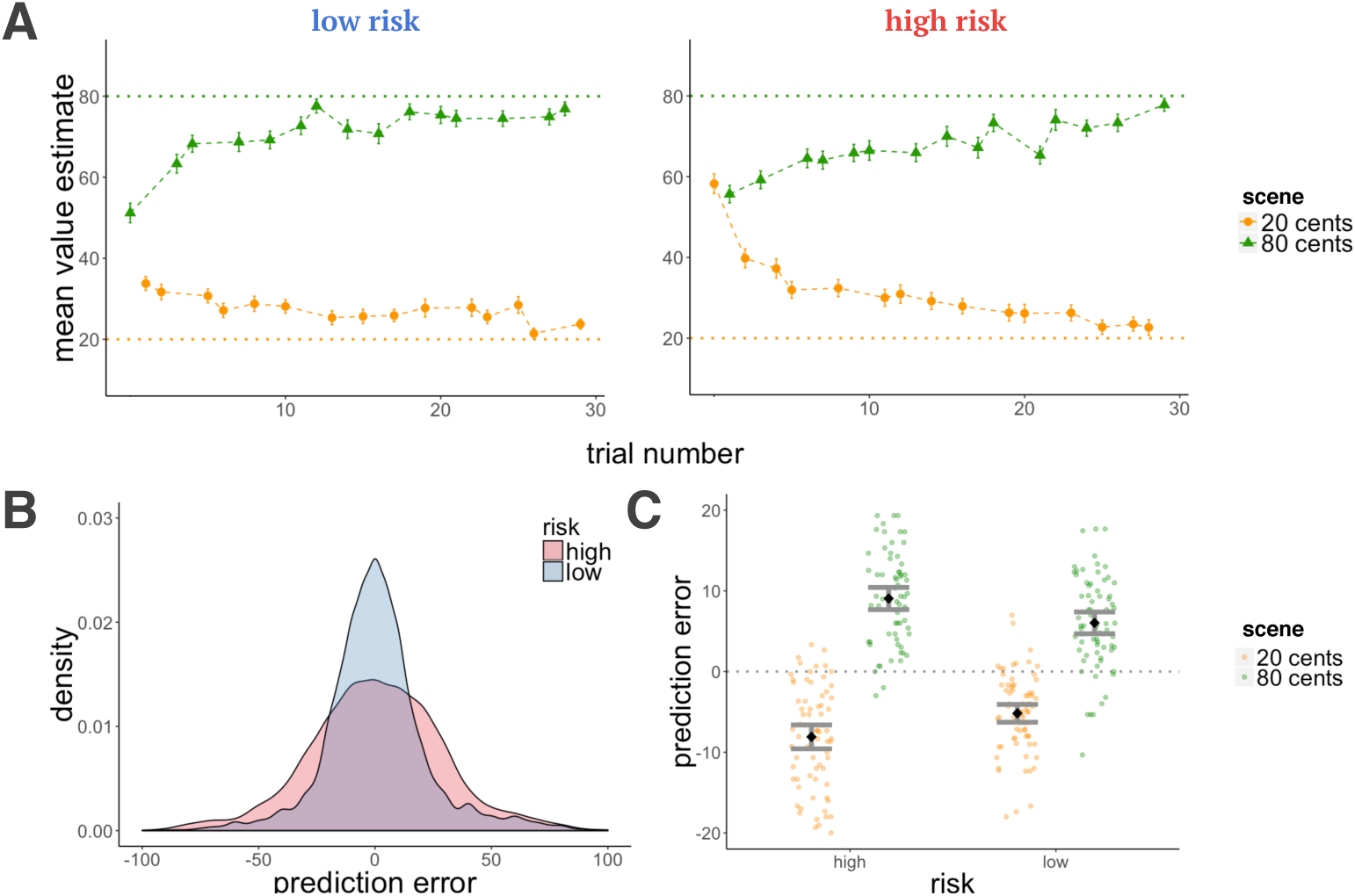
Experiment 3, learning results. A: Average estimates for the high and low-value scenes as a function of trial number for the high and low-risk rooms. Participants learned better in the low-risk room (although the difference in learning between risk rooms was more similar than in Exp. 1 & 2). B: Density plot of prediction errors in each room. There were more high-magnitude prediction errors in the low-risk room, making the range of prediction errors more similar between rooms than in Exp. 1 & 2. C: There was an interaction for prediction errors between risk context and scene value, such that participants overestimated the low-value scene and underestimated the high-value scene to a greater extent in the high-risk room. Error bars represent the standard error of the mean.

#### Memory by Risk and Prediction Error

We replicated the results of Experiments 1 and 2, and further found separate effects of context and unsigned prediction error on memory. A high-risk context and higher absolute prediction error enhanced memory for items, even with both predictors in the same model indicating independent effects (absolute prediction error: z = 2.24, p = 0.02, β = 0.12; risk: z = 2.58, p = 0.009, β = 0.24, Figure 11A-B). This effect was again significant when controlling for reward outcome (absolute prediction error: z = 2.72, p = 0.007, β = 0.15; reward: z = −0.38, p = n.s., β = −0.02) and value estimates (absolute prediction error: z = 2.70, p = 0.007, β = 0.15; value: z = −0.74, p = n.s., β = −0.03). We again found better sequence memory for high risk items, while controlling for the effect of distance (risk: z = 2.47, p = 0.01, β = 0.57; distance: z = 2.36, p = 0.02, β = 0.55). For source memory, we did not have the power to detect the effect in Experiment 2, and this difference was not statistically significant although it was in the same direction.

**Figure 11.**
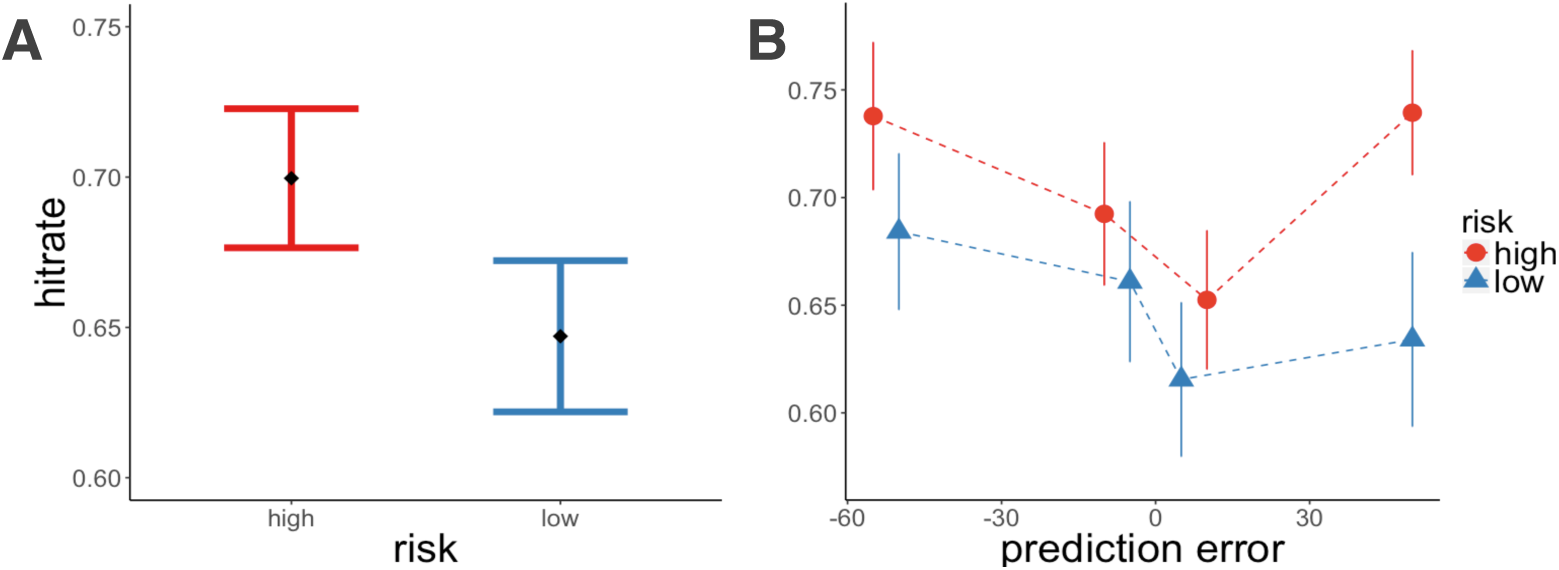
Experiment 3, recognition memory results. A: Recognition memory was better for high-risk items. B: Both absolute prediction errors and a high-risk context independently enhanced memory. Item memory was binned by the quartile values of prediction errors within each risk room. Each dot represents the average value within that quartile. Error bars represent the standard error of the mean.

It is worth noting here that there was a stronger effect of context in modulating memory than in Experiments 1 and 2 (the context effect remained when controlling for absolute prediction errors, unlike in Experiments 1 and 2). When learning between the two rooms was more similar, an independent effect of risk in increasing memory became apparent. One possible explanation for this finding is that memory-boosting effects of reward prediction errors might “spill over” to adjacent trials, enhancing memory for those items as well. To test for these “spill over” effects in the high-risk context, we measured whether previous and subsequent absolute prediction errors proactively or retroactively strengthened memory, while controlling for the absolute prediction error experienced on that trial. We did not find these effects for previous absolute prediction error (z = −1.71, p = n.s., β = −0.13) nor subsequent absolute prediction error (z = −0.93, p = n.s., β = −0.08), suggesting that this context effect may be due to general memory enhancement in a more arousing context.

#### Learning Rate by Risk and Prediction Error

As in Experiments 1 and 2, absolute prediction errors increased learning rates in both rooms, and there was a trend for higher learning rates in the low-risk room (absolute prediction error: t = 3.33, p < 0.001, β = 0.06; risk: t = 1.84, p = 0.06, β = 0.06; Figure 11A-B). We again did not find learning rate to predict memory on the current trial (z = −0.26, p = n.s., β = −0.01), nor the subsequent trial (z = −1.22, p = n.s., β = −0.08) while controlling for the effect of absolute prediction error on the current trial.

**Figure 12.**
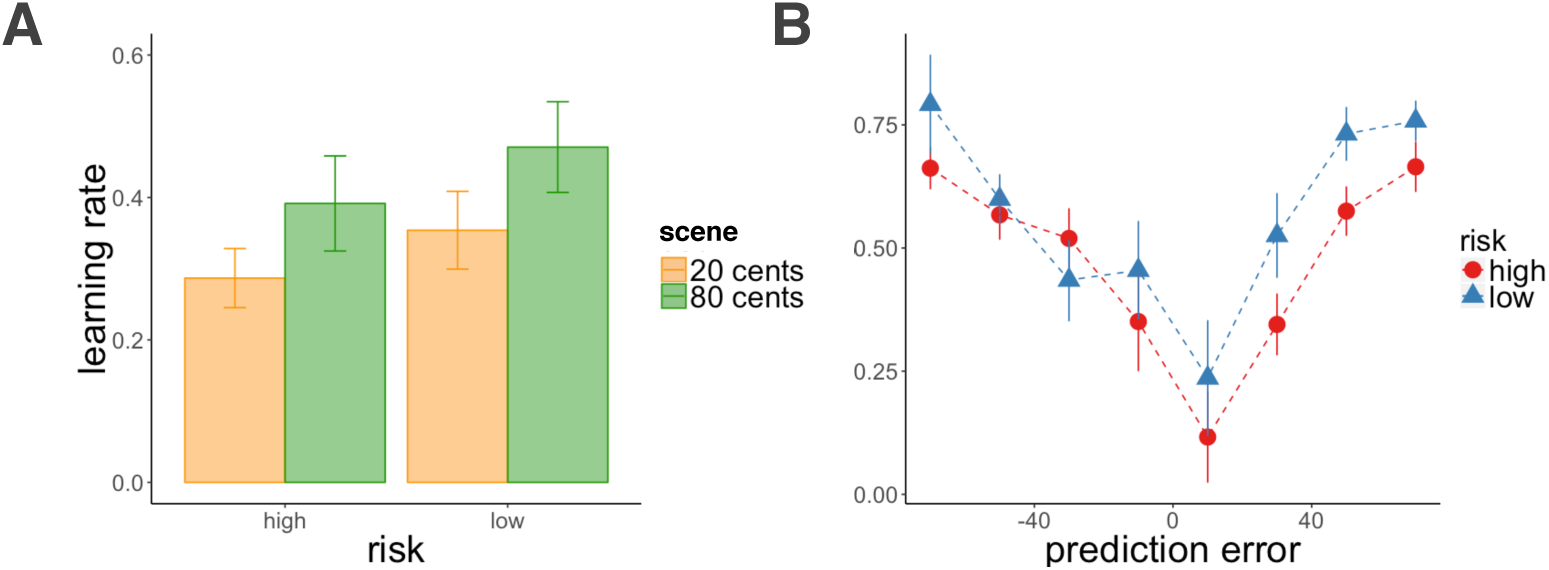
Experiment 3, learning rate results. A: There was a trend for higher average learning rate in the low-risk context. B: Absolute prediction errors increased learning rate. Learning rates were binned by prediction errors (each dot represents the average prediction error within the binned range). Error bars represent standard error of the mean.

#### Choice by Value Difference

As in Experiment 2, all choices (12 trials) were between images with matched reward outcomes. We replicated the results of Experiment 1 and 2, such that participants were more likely to choose the scene that they had initially valued more (z = 3.98, p < 0.001, β = 0.29, Figure 13).

**Figure 13.**
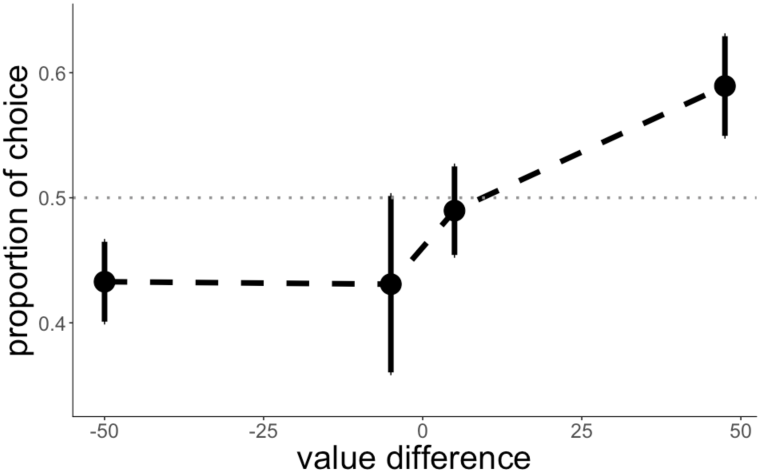
Experiment 3, choice results. For choices between options with matched reward outcomes, participants chose the scene that they had valued more. Each dot represents the average of the value differences binned by quartile. Error bars represent standard error of the mean.

## General Discussion

Our aim was to determine how reward prediction errors influence memory and decision making, above and beyond their known influence on learning. In Experiment 1, we demonstrated that unsigned, or absolute prediction errors enhanced memory for a rewarding event. That is, items that were accompanied by a large prediction error, whether positive (receiving much more reward than expected) or negative (receiving much less reward than expected) were better remembered in a subsequent test. We additionally found that risk context and absolute prediction errors modulated trial-by-trial learning rate – learning rate was higher in a low-risk environment, and for items that were accompanied by larger prediction errors. Notably, although unsigned prediction errors increased learning rate and enhanced memory, we did not find a trial-by-trial relationship between learning rate and memory. Moreover, a higher risk context led to lower learning rates but better memory on average, emphasizing distinct underlying mechanisms.

In Experiment 2 we allowed for more learning in each room and replicated all the effects from Experiment 1, while further demonstrating that a high-risk context improved source and sequence memory. In Experiment 3, we set out to test whether risk per se affected learning and memory, or if our findings could be attributed to the higher overlap between the reward distributions of the two options in the high-risk context. For this, we eliminated the overlap in outcome distributions in both contexts, and reproduced the original results. This manipulation resulted in a more similar range of prediction errors in both risk contexts, which allowed us to demonstrate an effect of context on episodic memory that was separate from the effect of absolute prediction errors.

Previous work has shown both a collaboration between learning and memory systems, such as boosting of memory for items experienced during reward anticipation (Adcock et al., 2006) including oddball events (Murty & Adcock, 2014), as well as a competition between the systems, where the successful encoding of items experienced prior to reward outcome is thought to interfere with striatal prediction errors (Wimmer, Braun, Daw, & Shohamy, 2014). Here we showed how learning mechanisms, namely absolute prediction errors, impact episodic memory for the actual rewarding event, similar to the enhanced encoding of unexpected paired associates (Greve, Cooper, Kaula, Anderson, & Henson, 2017).

An effect of (signed) dopaminergic prediction errors from the VTA to the hippocampus would have predicted an asymmetric effect on memory, such that memories benefit from a positive prediction error (signaled by an increase in dopaminergic firing from the VTA), but not a negative prediction error (signaled by a dip in dopaminergic firing). Instead, we found that the absolute magnitude of prediction errors, regardless of the sign, affected memory. This mechanism perhaps explains the finding that extreme outcomes are recalled first, are perceived as having occurred more frequently, and increase preference for a risky option (Ludvig, Madan, & Spetch, 2014; Madan, Ludvig, & Spetch, 2014).

The mnemonic effects of unsigned prediction errors, likened to surprise and contributing to arousal, have been linked to the noradrenergic locus coeruleus, and are thought to bias encoding of relevant or “high priority” stimuli (Clewett, Schoeke, & Mather, 2014; Mather, Clewett, Sakaki, & Harley, 2015; Miendlarzewska, Bavelier, & Schwartz, 2016). Our finding that absolute prediction errors influenced subsequent memory fits with the mechanism described in the *Introduction*, whereby the locus coeruleus-norepinephrine (LC-NE) system responds to salient (surprising) events, and dopamine co-released with norepinephrine in the locus coeruleus strengthens hippocampal memories (Kempadoo et al., 2016; Takeuchi et al., 2016). This proposed mechanism would seem to imply that increases in learning rate (previously linked to norepinephrine release) and enhanced memory (linked to dopamine release) should be correlated across trials, given the hypothesized common cause of LC activation.

However, we found that – across trials – increases in learning rate were uncorrelated with enhanced memory, suggesting that the actual mechanism may involve additional (or different) steps from the one described above.

In our task, learning rate not only increased with the magnitude of prediction error, but also changed with the risk of the environment. In line with our results, recent work shows that learning rate scales to reward variance, with higher learning rates in lower variance contexts (Diederen & Schultz, 2015; Diederen et al., 2016). Greater sensitivity to the same magnitude prediction errors in a low-versus a high-variance environment demonstrates adaptation to reward context, where in a low-risk context, even small prediction errors are more relevant to learning than they would be when there is greater reward variance. This heightened sensitivity to changes in the low-risk environment, however, was not associated with improved episodic memory in our experiments. In fact, in Experiment 3, we found that memory was better for *high-risk* items, even when controlling for trial-by-trial variance in reward prediction error. The opposing effects of risk on learning rate and episodic memory again suggest distinct underlying mechanisms, in agreement with work characterizing learning and memory systems as separate and even antagonistic given the task at hand (Foerde, Braun, & Shohamy, 2012; Wimmer et al., 2014).

To explain the beneficial effect of high-risk environments on episodic memory, we hypothesized that better memory for high prediction error events could potentially “spill over” to surrounding items, in line with work showing that inducing an “encoding” state (such as through the presentation of novel items) introduces a lingering bias to encode subsequent items (Duncan & Shohamy, 2016; Duncan et al., 2012). These effects, however, did not explain how risk context modulated memory in our task, as we did not find prediction error events to additionally improve memory for adjacent items. Instead, we attribute this context effect to improved encoding when in a more aroused state, although future studies should more directly characterize the link between arousal and enhanced memory in risky environments.

Finally, when given a choice, people preferred scenes that had been previously associated with greater reward outcomes. Surprisingly, however, when choosing between images that had the same reward outcomes, people preferred the ones they had initially guessed would lead to a higher reward. This perhaps points to anticipatory dopaminergic activation at image onset (Adcock et al., 2006) influencing later preference, regardless of the actual outcome (and its associated subsequent disappointment). In particular, we did not find an effect of absolute prediction error or risk context on choice, which could be due to the statistical power of our task in detecting an effect, or could demonstrate that item memory may not necessarily guide value-based choice. This latter point is echoed in a recent study showing that memory for the values associated with items, as opposed to just item recognition, predicts value-based decision making, which highlights the role of enriched episodic memory (instead of decontextualized recognition) in supporting adaptive decision making (Murty, FeldmanHall, Hunter, Phelps, & Davachi, 2016). Even though we did not find absolute prediction error or risk context effects on later choice preference, it remains to be determined whether memories enhanced by large prediction errors may still bias decision making by prioritizing which experiences are sampled or reinstated during value-based decision making.

In conclusion, we show that surprising rewards and high-risk contexts improve memory, revealing that error and risk modulate memory in addition to the mnemonic effects of high rewards previously reported in the literature. We further demonstrated that absolute prediction errors, associated with locus coeruleus modulation, have dissociable effects on learning rate and memory, pointing to separate influences on incremental learning and episodic memory.

## Acknowledgments

This work was supported by the Ellison Foundation (Y.N.), grant R01MH098861 from the National Institute for Mental Health (Y.N.), grant W911NF-14-1-0101 from the Army Research Office (Y.N.), and the National Science Foundation’s Graduate Research Fellowship Program (N.R.).

